# The color pattern inducing gene *wingless* is expressed in specific cell types of campaniform sensilla of a polka-dotted fruit fly, *Drosophila guttifera*

**DOI:** 10.1101/2021.02.01.429284

**Authors:** Masato Koseki, Nobuaki K. Tanaka, Shigeyuki Koshikawa

## Abstract

A polka-dotted fruit fly, *Drosophila guttifera,* has a unique pigmentation pattern on its wings and is used as a model for evo-devo studies exploring the mechanism of evolutionary gain of novel traits. In this species, a morphogen-encoding gene, *wingless*, is expressed in species-specific positions and induces a unique pigmentation pattern. To produce some of the pigmentation spots on wing veins, *wingless* is thought to be expressed in developing campaniform sensilla cells, but it was unknown which of the four cell types there express(es) *wingless*. Here we show that two of the cell types, dome cells and socket cells, express *wingless*, as indicated by *in situ* hybridization together with immunohistochemistry. This is a unique case in which non-neuronal SOP (sensory organ precursor) progeny cells produce Wingless as an inducer of pigmentation pattern formation. Our finding opens a path to clarifying the mechanism of evolutionary gain of a unique *wingless* expression pattern by analyzing gene regulation in dome cells and socket cells.

## Introduction

Animal color patterns are an example of the morphological diversity of organisms. Ecological roles of color patterns have been studied (Cott 1940), and another major issue regarding color patterns is their formation process. In particular, color pattern formation of insects has been investigated to gain insight into the relationship between regulation of gene expression and morphological evolution. As a result of studies to elucidate the process of pattern formation, patterning genes whose expression induces the color formation have been identified. For example, the pattern of *optix* expression determines the position of red pigmentation in adult wings of *Heliconius* butterflies (Reed et al. 2011; Martin et al. 2014; Zhang et al. 2017), *WntA* expression determines the border of pigmentation in adult wings of Nymphalidae butterflies (Martin et al. 2012; Martin and Reed 2014; Mazo-Vargas et al. 2017), and *pannier* expression determines the aposematic color pattern in the adult ladybird beetles (Ando et al. 2018; Gautier et al. 2018). Interestingly, these genes are also expressed during ontogenesis. This indicates an evolutionary process in which genes with roles in ontogenesis were co-opted for color pattern formation (Jiggins et al. 2017). In order to examine in detail the evolutionary process that produces color patterns, it is necessary to elucidate the mechanism of spatiotemporal regulation of patterning genes (Fukutomi and Koshikawa 2021).

Various pigmentation patterns are found in adult wings of *Drosophila* fruit flies (Insecta, Diptera, Drosophilidae) (Wittkopp et al. 2002; Massey and Wittkopp 2016; Dufour et al. 2020; Koshikawa 2020; Werner et al. 2020). Adult *Drosophila guttifera* flies have a species-specific wing spot pattern (Fig. 1A). The spots occur at specific positions, such as around campaniform sensilla on wing veins (Werner et al. 2010; Koshikawa et al. 2015; Fukutomi et al. 2017; Fukutomi et al. 2021). The spot formation around campaniform sensilla is a suitable model for examining the spatiotemporal regulation of the patterning gene (Koshikawa et al. 2017). This formation is induced by the species-specific expression of the patterning gene *wingless* (*wg*) during the pupal stage (Fig. 1C; Werner et al. 2010; Koshikawa et al. 2015). *wg* is expressed at campaniform sensilla on wing veins of *D. guttifera* (Fig 1C), and this expression is not detected in other *Drosophila* species (Koshikawa et al. 2015). The expression begins at mid-pupa (late stage 6) and then induces the subsequent expression of pigmentation genes such as *yellow* (Werner et al. 2010). In *D. melanogaster*, Each campaniform sensillum, which is a mechanoreceptor involved in flight control by sensing wing flexion (Tuthill and Wilson 2016), consists of four differentiated cells: a socket (tormogen) cell, a dome (trichogen) cell, a sheath (thecogen) cell, and a neuron (Fig. 1B; Van De Bor et al. 2000; Van De Bor et al. 2001). The formation of a campaniform sensillum resembles that of a bristle, a type of mechanoreceptor. The sensory organ precursor (SOP) selected from a proneural cluster (Furman and Bukharina 2008; Gómez-Skarmeta et al. 1995) divides into the two secondary precursors, PⅡa and PⅡb (Van De Bor et al. 2000; Van De Bor and Giangrande 2001). The socket and dome cells are generated by the division of the PⅡa, and the sheath cell and neuron are generated by the division of the tertiary precursor PⅢb, which is a progeny of the PⅡb (Van De Bor et al. 2000; Van De Bor and Giangrande 2001). Subsequently, the differentiation process of each of these cells follows (Furman and Bukharina 2008), but the detailed gene expression profiles involved in the differentiation have not been well investigated. The expression of the patterning gene *wg* begins during the differentiation stage of the four cells of the campaniform sensilla in *D. guttifera* (Werner et al. 2010), indicating that the cell differentiation and *wg* expression are synchronized. Understanding the relationships between campaniform sensilla differentiation and *wg* expression is a key for unravelling the process of evolutionary gain of species-specific *wg* expression.

**Figure 1.**
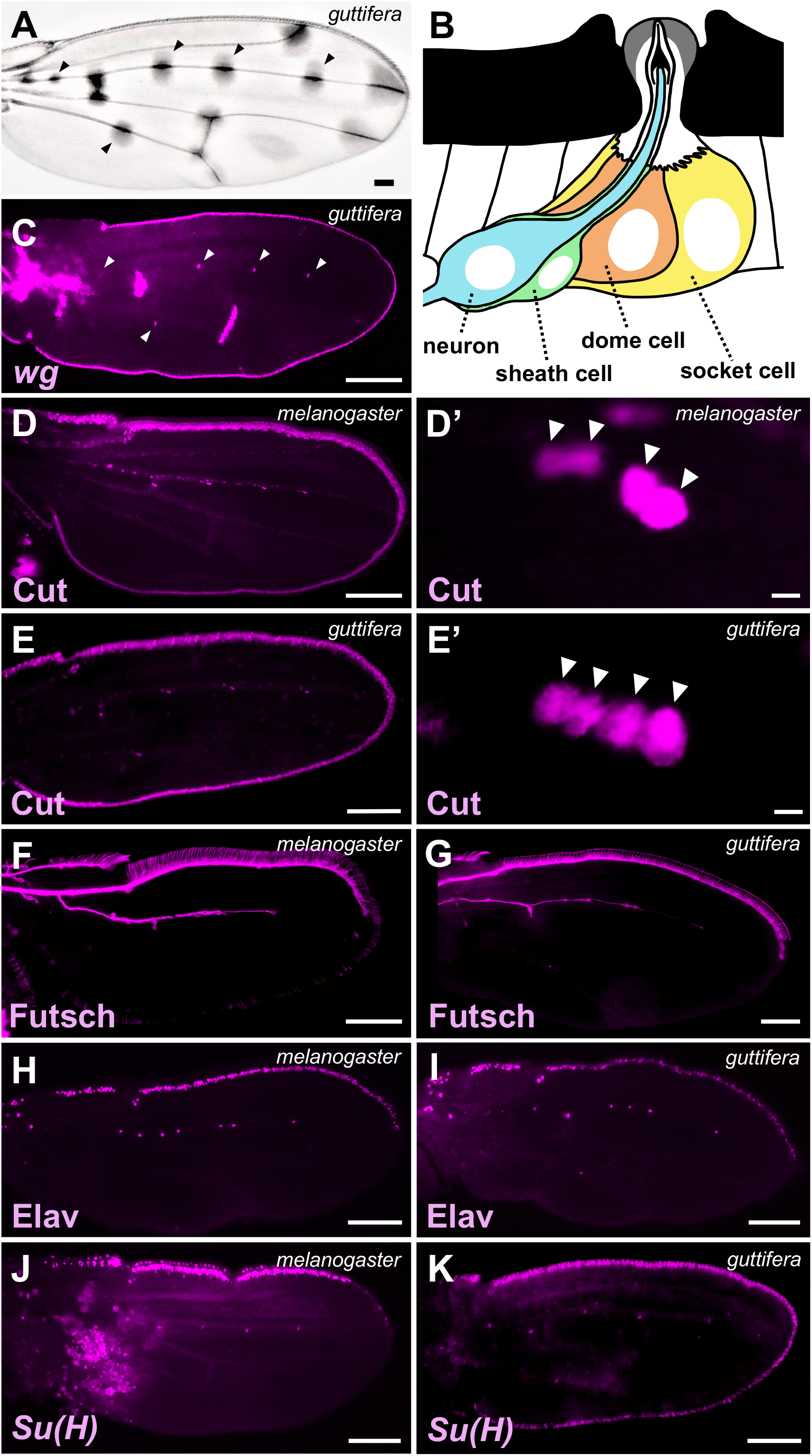
Expression patterns of *wingless* (*wg*) and developmental marker genes in pupal wings. Transcripts were visualized by *in situ* hybridization using Fast Red, and proteins were visualized by immunostaining with Alexa555. **A** An adult wing of *D. guttifera*; arrowheads indicate the pigmentation around campaniform sensilla. **Bn** Transverse section diagram of campaniform sensillum (After Gullan and Cranston, 2014). **C** Expression pattern of the patterning gene *wg* in a pupal wing of *D. guttifera*; arrowheads indicate the expression at campaniform sensilla. **D and D’** Localization of Cut protein in *D. melanogaster.* **E and E’** Localization of Cut protein in *D. guttifera.* Cut was observed in nuclei of all cells composing campaniform sensilla in both species. Arrowheads indicate the nuclei. **F** Localization of Futsch protein in *D. melanogaster*. **G** Localization of Futsch protein in *D. guttifera*. Futsch was observed in cytoplasm of neurons in both species. **H** Localization of Elav (Embryonic lethal abnormal vision) protein in *D. melanogaster.* **I** Localization of Elav protein in *D. guttifera.* Elav protein was observed in nuclei of neurons in both species. **J** Localization of *Suppressor of Hairless* [*Su(H)*] transcripts in *D. melanogaster*. **K** Localization of *Su(H)* transcripts in *D. guttifera*. *Su(H)* transcripts were observed in socket cells in both species. Scale bar indicates 100 μm (**A, B, D-K**) or 5 μm (**D’, E’**). Distal is to the right.

Proteins of the Wnt family, including Wingless (Wg) protein, are involved in the formation of structures characteristic of the neuronal network, such as synapses and axons (Packard et al. 2002; He et al. 2018). Considering that *wg* may be expressed in neurons in general, although *wg* is not expressed at campaniform sensilla of *D. melanogaster*, it is possible that co-option of *wg* expression occurs in neurons of campaniform sensilla of *D. guttifera*. Here, as the first step to elucidate the relationships between the differentiation of cells composing campaniform sensilla and the pigmentated spot formation, we investigated whether neurons express the patterning gene *wg*, and if not, which cells express *wg*. The identification was performed by the dual detection of *wg* transcripts by *in situ* hybridization and of specific marker protein or cell membrane of campaniform sensilla by immunohistological staining in pupal wings.

## Materials and methods

### Flies and genomic DNA

We used *D. melanogaster* Oregon-R (wild-type) and *D. guttifera* (stock no. 15130-1971.10, obtained from the *Drosophila* Species Stock Center at the University of California, San Diego), for genomic DNA preparation and gene expression analysis. Both fly lines were reared on standard food containing cornmeal, sugar, yeast, and agar at room temperature (Fukutomi et al. 2018).

### Dissection and Fixation

Dissection of pupal wings was performed as described previously (Werner et al. 2010). After the pupal membrane was removed, pupal wings were fixed in PBS (phosphate buffered saline, Takara Bio) with 4% paraformaldehyde (PFA) for 20 minutes (min) at room temperature. Fixed samples were washed three times in PBT (0.1% Triton X-100 in PBS) and stored in methanol at −20℃.

### Immunohistochemistry

Stored wing samples were incubated with primary antibodies in PBT overnight at 4℃. Following three washes with PBT, samples were incubated with fluorescent secondary antibodies in PBT for 2 h at room temperature and washed three times with PBT. After the last PBT wash, samples were mounted with Vectashield Mounting Medium with DAPI (Vector Laboratories). Each wash was 5 min long. The following primary antibodies were used at the indicated dilutions: mouse anti-Cut, 1:1000 [2B10; Developmental Studies Hybridoma Bank (DSHB)]; mouse anti-Elav (Embryonic lethal abnormal vision), 1:1000 (9F8A9, DSHB); mouse anti-Futsch, 1:1000 (22C10; DSHB); mouse anti-Na^+^/K^+^-ATPaseα (Chicken homolog of *Drosophila* Atpα), 1:50 (a5; DSHB). For secondary antibodies, anti-mouse-Alexa555 conjugate (Abcam) or anti-mouse-Alexa488 conjugate (Abcam) was used at 1:500.

### *In situ* hybridization

Digoxygenin-labeled antisense RNA probes of *wg* and *Suppressor of Hairless* [*Su(H)*] were produced as described previously (Werner et al. 2010). Genomic DNA was extracted by using a DNeasy Blood & Tissue Kit (QIAGEN). The following forward and reverse primers were used to amplify a 377 bp DNA fragment of *wg* exon 2 in *D. guttifera*: 5’-CACGTTCAGGCGGAGATGCG-3’ and 5’-GGCGATGGCATATTGGGATGATG-3’, a 525 bp DNA fragment of *Su(H)* exon 2 in *D. guttifera*: 5’-CAGTGATCAGGATATGCAGC-3’ and 5’-TGCGAAACAGGATCATCAGC-3’, and a 402 bp DNA fragment of *Su(H)* exon 2 in *D. melanogaster*: 5’-AGCTGGATCTCAATGGCAAG-3’ and 5’-CATTCATTACGGAGCCACAG-3’. DNA fragments were cloned into pGEM-T Easy Vector (Promega). DNA templates for *in vitro* transcription were amplified using M13F and M13R primers. RNA probes were transcribed *in vitro* with T7 or SP6 polymerase (Promega) and DIG (Digoxigenin) RNA Labeling Mix (Roche). Each probe was purified using a ProbeQuant G-50 Micro Column (Cytiva) and stored in RNase-free water at −20℃. Stored wing samples were treated with methanol containing 2% H₂O₂ for 20 min at room temperature, as described previously (Lauter et al. 2011). Then they were washed twice with ethanol, incubated in a mixture of xylene and ethanol (1:1 v/v) for 60 min, washed three times with ethanol, and rehydrated by two washes with methanol and two washes with PBT. After treatment with a mixture of acetone and PBT (4:1 v/v) for 14 min at −20℃, samples were washed twice with PBT, post-fixed in PBS with 4% PFA for 20 min and washed three times with PBT. The fixation in acetone was performed with reference to Nagaso et al. (2001). The hybridization process and anti-DIG antibody incubation were performed as described previously (Sturtevant et al. 1993, Werner et al. 2010), except that the hybridization buffer contained 5% dextran sulfate and the hybridization temperature was 57℃. Signals of transcripts were detected using Anti-DIG-AP (alkaline phosphatase) and Fab fragments from sheep (Roche) and developed in Fast Red TR and naphthol-AS-MX-phosphate in 0.1 M Tris-HCl pH 8.2 (tablet set; Sigma). After three washes with PBT, samples were mounted in PBT or 50% glycerol diluted with PBS.

### Dual detection

The pretreatment and hybridization processes were performed as described above. After washes with PBT, hybridized wings were incubated with Anti-DIG-AP Fab fragments (Roche) diluted 1:6000 and primary antibodies in Pierce Immunostain Enhancer (PIE) (Thermo Scientific) overnight at 4℃. Then they were washed three times with PBT, incubated with fluorescent secondary antibodies in PIE for 2 h at room temperature and washed three times with PBT. Detection of transcripts, subsequent washes, and mounting were performed as described in the “*In situ* hybridization” section.

### Microscopy and image analysis

Preparations were observed using a BX60 microscope (Olympus) or an LSM700 confocal microscope (Carl Zeiss). Confocal images were taken at z–intervals of 0.4 μm. The brightness, contrast, and color of images were adjusted with Fiji (Schindelin et al. 2012).

## Results and discussion

### Comparison of gene expression patterns in pupal wings of *D. melanogaster* and *D. guttifera*

We first investigated whether *cut*, *futsch*, *embryonic lethal abnormal vision* (*elav*), and *Suppressor of Hairless* [*Su(H)*] are expressed in the campaniform sensilla on pupal wings of *D. guttifera* (Fig. 1). These are marker genes expressed in the cells of sensilla in *D. melanogaster.* Cut protein (Blochlinger et al. 1990; 1993) is localized in the nuclei of all four types of cells composing campaniform sensilla during the developmental stages in *D. melanogaster* (Van De Vor et al. 2000). Futsch, also known as the antigen of 22C10 antibodies, and Elav proteins are localized in the cytoplasm and nucleus of neurons, respectively (Fig. 1F, H; Hummel et al. 2000; Aigouy et al. 2004; Robinow and White 1988; Van De Bor et al. 2000). *Su(H)* is a gene specifically expressed in the socket cells composing bristles of *D. melanogaster* (Barolo et al. 2000).

In *D. melanogaster*, we found that Cut expression was observed in the sensilla of the wing margin, the third longitudinal vein, and the anterior cross vein of pupal wings (Fig. 1D). The magnified image of L3-1 (Fig. 1D’) shows four cells with a high level of accumulation of Cut, as in other campaniform sensilla of the third longitudinal vein (L3-2 and L3-3). In *D. guttifera*, Cut was localized in the campaniform sensillum on the fifth longitudinal vein (L5) in addition to the third longitudinal vein (Fig. 1E). This is in accord with the previous report that there is a campaniform sensillum on the fifth longitudinal vein in the wing of *D. guttifera* (Sturtevant 1921), but not in *D. melanogaster*. In a magnified view of the *D. guttifera* wing (L3-1, Fig 1E’), four cells containing a high level of Cut protein are seen, as in *D. melanogaster*, indicating that a campaniform sensillum of *D. guttifera* also consists of four types of cells.

Futsch protein, also known as 22C10 antigen, was detected throughout the cytoplasm of neurons in *D. melanogaster* (Fig. 1F; Hummel et al. 2000; Aigouy et al. 2004). A similar pattern of Futsch localization in pupal wings was observed in *D. guttifera* (Fig. 1G). Elav protein is localized in nuclei of neurons of *D. melanogaster* (Fig. 1H; Robinow and White 1988; Van De Bor et al. 2000). A similar pattern of Elav localization in pupal wings was observed in *D. guttifera* (Fig. 1I), suggesting that Elav is localized in nuclei of neurons of *D. guttifera*. In summary, there was no substantial difference in the morphology or position of neurons in pupal wings between the two species.

The expression of *Su(H)* was detected at the estimated position of the campaniform sensilla in *D. melanogaster* (Fig. 1J), suggesting that the socket cells composing the campaniform sensilla of *D. melanogaster* express *Su(H)*. Similarly, the expression of *Su(H)* was observed in the campaniform sensilla of *D. guttifera* (Fig. 1K). This indicates that the position of socket cells in pupal wings of *D. guttifera* can be visualized by detecting the expression of *Su(H)*. However, it was unknown whether the detection of *Su(H)* could visualize the entire area of the socket cells.

### Neurons and sheath cells do not express the patterning gene *wg*

Next, we analyzed whether *wg* is expressed in the neurons composing campaniform sensilla in the pupal wings of *D. guttifera*. We simultaneously detected *wg* transcripts and the neuronal marker gene expression in the four sensilla, L3-1, L3-2, L3-3, and L5 (Fig 2), where *wg* is expressed at a high level. We found that the signals of Elav protein localized in neuronal nuclei (Fig. 2A-D) and *wg* transcripts (Fig. 2A’-D’) were not colocalized within the four campaniform sensilla (Fig. 2A”-D”). In addition, we did not observe colocalization of Futsch protein, which was localized throughout the neuronal cytoplasm (Fig. 2E-H), and *wg* transcripts (Fig. 2E’-H’) in the four campaniform sensilla (Fig. 2E”-H”). These results indicate that the patterning gene *wg* is not expressed in the neurons in the campaniform sensilla.

**Figure 2.**
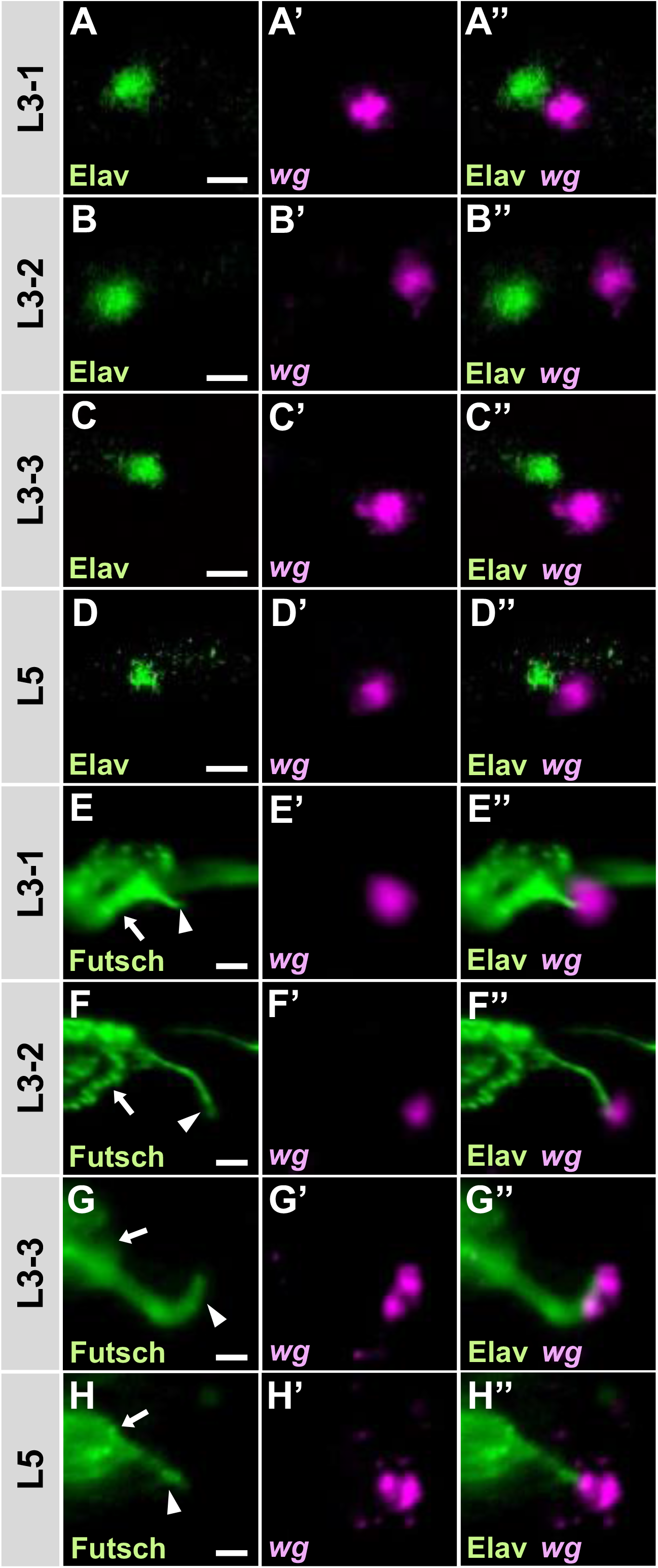
The patterning gene *wingless* (*wg*) is not expressed in neurons of campaniform sensilla. Signals of Elav (Embryonic lethal abnormal vision) protein (**A**, **B**, **C**, **D**) localized in neuronal nuclei were compared with those of *wg* transcripts (**A’**, **B’**, **C’**, **D’**). Signals of Elav did not overlap with those of *wg* (**A’’**, **B’’**, **C’’**, **D’’**). Signals of Futsch protein (**E**, **F**, **G**, **H**) localized in neuronal cytoplasm compared with those of *wg* transcripts (**E’**, **F’**, **G’**, **H’**). Arrows indicate the cell bodies and arrowheads indicate the dendrites. Signals of Futsch did not overlap with signals of *wg* (**E’’**, **F’’**, **G’’**, **H’’**). Scale bar indicates 5 μm. Distal is to the right. Elav and Futsch proteins were visualized by immunostaining with Alexa488. *wg* transcripts were visualized by *in situ* hybridization using Fast Red.

The signal of *wg* transcripts was observed in contact with the tip of the dendrite labeled with anti-Futsch antibody at the level of detection by light microscopy (Fig. 2E’’-H’’). With reference to the structure of a campaniform sensillum (Fig. 1B; Chevalier 1969; Gullan and Cranston 2014), the relatively large socket and dome cells surround the dendrite. In contrast, the sheath cell covers the neuron from the inner dendrite to the cell body and appears not to spread around the tip of the dendrite. In addition, *wg* transcripts appeared to be localized in two spots in some images (Fig. 2G’, H’), indicating the possibility that two cells express *wg*. The *wg* expression pattern suggests that *wg* is expressed not in sheath cells but in either socket cells or dome cells, or both.

### Both socket and dome cells express the patterning gene *wg*

In order to determine whether socket and/or dome cells composing campaniform sensilla express *wg*, we furthermore analyzed the expression pattern of *wg*. As cytoplasmic markers of sheath, socket and dome cells have not been identified in *D. guttifera*, we labeled the campaniform sensilla with anti-Na^+^/K^+^-ATPase α subunit (Atpα) antibody (Lebovitz et al. 1989), a marker of the cell membranes. The signals visualized three concentric layers within the sensilla (Fig 3A). To identify the cell type of each layer, we simultaneously detected *Su(H)* transcripts, which are expressed in the socket cells in the mechanoreceptor bristles. We found that *Su(H)* signals (Fig. 3B’, C’) were limited to the outermost layer (Fig 3B”, C”), indicating that the outermost layer is composed of the socket cell, as shown in Fig 1B. Although we have not identified the cell types of all of the three layers with specific markers, based on the cell arrangement shown in Fig 1B, we conclude that the innermost layer is the neuronal dendrite, the second most is the dome cell, and the outermost is the socket cell. Among these three concentric layers, we found that *wg* is expressed in the two outer layers, both socket and dome cells (3D”, D’”, E”, E”’).

**Figure 3.**
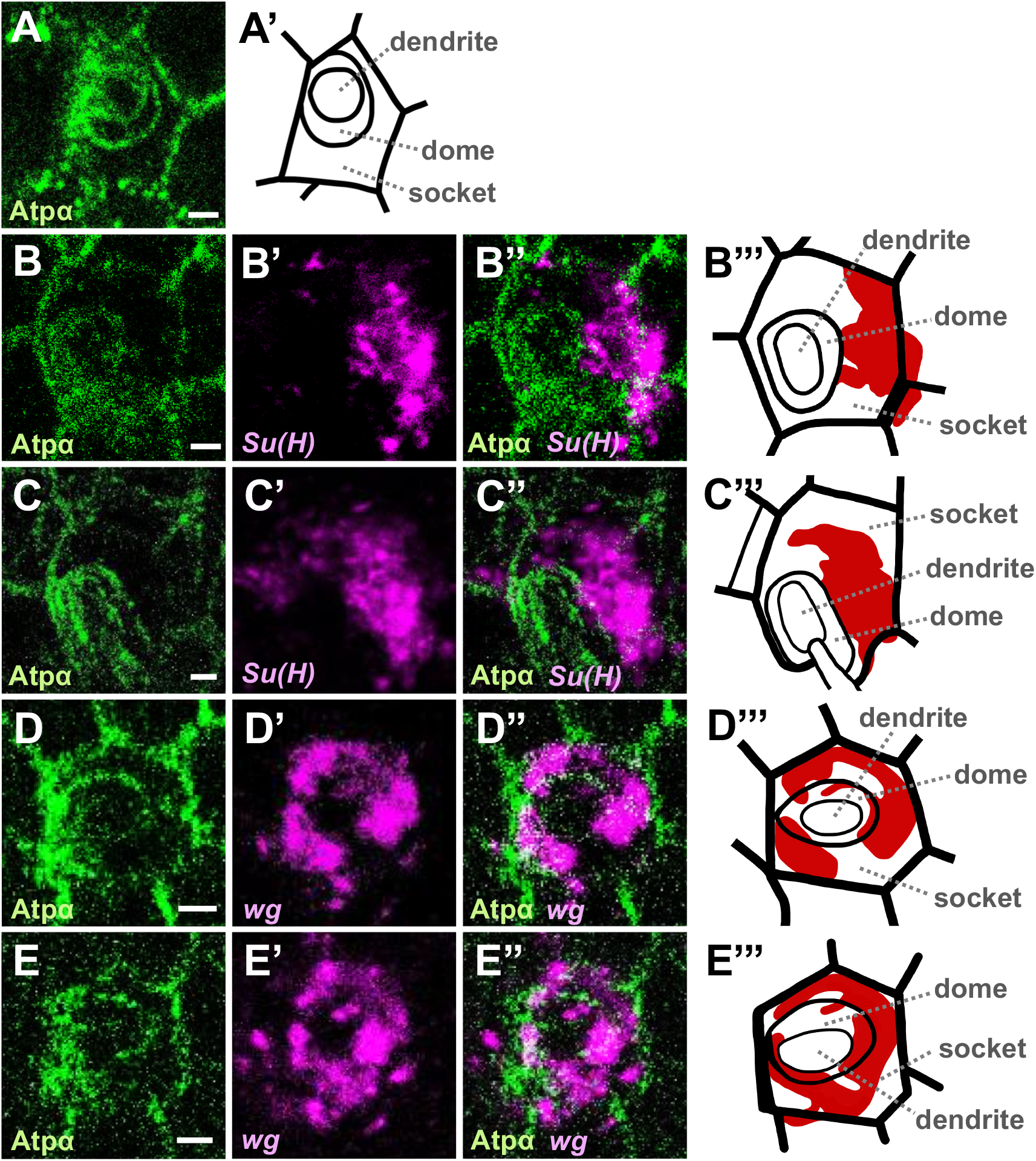
*wg* transcripts were observed in the socket and dome cells. The signal of Na+/K+-ATPaseα (Atpα, green in the figures) visualizes the cell membrane at campaniform sensilla (**A**, **B**, **C**, **D**). *Suppressor of Hairless* [*Su(H)*, magenta in the second left and third left figures] transcripts were detected at campaniform sensilla (**B’**, **C’**). Cell types can be identified according to their morphology and relative position. The relatively large cell surrounding the other structures of the campaniform sensillum and expressing *Su(H)* transcripts was identified as a socket cell (**A’, B’’**, **B’’’, C’’**, **C’’’**). The cell surrounded by a socket cell was identified as a dome cell. The sheath cell was not obvious in this plane, while the dendrite surrounded by a sheath cell was visible. *wg* transcripts (**C’**, **D’**) were localized in the socket cell and the inner dome cell (**C’**’, **C’’’**, **D’’**, **D’’’**). All panels show L3-1 sensillum. Scale bar indicates 1 μm. Atpα protein was visualized by immunostaining with Alexa488. *wg* and *Su(H)* transcripts were visualized by *in situ* hybridization using Fast Red.

This study revealed that *wg* expression which induces pigmentation spots around campaniform sensilla of *D. guttifera* occurs in the socket and dome cells, but not neurons. Considering that Wg protein is known to act as a morphogen, it is expected that Wg produced by these two cells is secreted to the surrounding epidermal cells and induces expression of the pigmentation genes in the recipient cells. This process of color pattern formation, in which co-option of the patterning gene occurs in specific cells composing nerve tissues, has not been reported before.

Although it is still unclear what gene regulatory network is responsible for the co-option of *wg* in these two cell types, a clue was obtained from an abnormal individual of *D. guttifera* (Werner et al. 2010). In the abnormal individual, the dome structure of a campaniform sensillum in the adult wing was converted to the bristle structure, and the pigmentation spot was not formed around the structure. Considering our findings, suppression of the *wg* expression by the fate change of socket and/or dome cells can be assumed to have been the reason for the abnormality of pigmentation in that individual. This suggests that we can approach the regulatory process of the spatiotemporal expression of *wg* by clarifying the state of gene expression underlying the fate change of these two cells. Furthermore, in *D. melanogaster*, overexpression of the *hindsight* gene in sensory organ precursors (SOPs) on the future wing vein of the wing disc transformed the dome structure of some campaniform sensilla to the bristle-like structure (Szablewski and Reed 2019). The expression of *hindsight* was confirmed in SOPs (Buffin and Gho 2010), and *hindsight* encodes a transcription factor that is necessary to positively regulate EGFR signaling (Kim et al. 2020). If the transformation mechanism applies to the reason why the described abnormality of *D. guttifera* occurred, the expression of *wg* could be under the control of EGFR signaling, and overactivation of the signaling could downregulate the *wg* expression. In order to confirm whether EGFR signaling influences the spatiotemporal expression of *wg* in socket and dome cells, it will be necessary to investigate whether the spot formation of *D. guttifera* is influenced by manipulating the expression of genes involved in EGFR signaling, such as *hindsight*.

## Acknowledgements

We thank Elizabeth Nakajima for English editing, and Yuichi Fukutomi and Tomohiro Yanone for technical advice.

## Funding

This work was supported by MEXT/JSPS KAKENHI (18H02486) grant to S. K.

## Notes

### Competing Interest Statement

The authors have declared no competing interest.

